# Deep behavioural phenotyping reveals divergent trajectories of ageing and quantifies health state in *C. elegans*

**DOI:** 10.1101/555847

**Authors:** Céline N. Martineau, Bora Baskaner, Renée I. Seinstra, William R. Schafer, André E. X. Brown, Ellen A. A. Nollen, Patrick Laurent

## Abstract

Neurodegenerative diseases may be the cause or the consequence of an acceleration of physiological ageing. Evidence for this concept is lacking due to practical limitations of human studies. Here, we compared the processes of physiological and pathological ageing of individual *C. elegans* over their lifespan. Using multi-parametric phenotyping, trajectories of ageing can be defined within a phenotypic landscape made of a large set of phenotypical features. Rather than an acceleration of ageing, a model for synucleinopathy showed a divergent trajectory of ageing. The pathological progression in individual animals can be predicted from early phenotypes with high accuracy. Despite of similar lifespans, disease-model worms display an early onset of decline in their phenotypic range of ability. This loss of flexibility provides an index of health valid for physiological and pathological contexts. Finally, we demonstrate the power of multi-parametric dataset to describe ageing, to quantify health and to predict specific health risks.

## Introduction

Ageing is an important factor affecting quality of life and is a major risk factor for diseases. Because of its short lifespan, the nematode *Caenorhabditis elegans* has been used for decades as a model organism to study normal ageing and to model age-related disorders like neurodegenerative diseases. The measurement of survival curves has been a key parameter to identify molecular pathways modulating longevity ^1,2^. Importantly, the survival curves merely quantify the very end of the complex health state progression. Therefore, efforts have been made to define health parameters to better describe *C. elegans* ageing ^3–5^. *C. elegans* ageing is marked by morphological and functional changes ^6–10^. Recently, a limited number of these biomarkers of age have been used for a longitudinal description of the physiological ageing process in a closed system ^5^.

Ideally suited for longitudinal studies, the morphological and behavioural repertoire of the worm offers numerous easily quantifiable non-invasive parameters. Importantly, the behaviour of *C. elegans* evolves from the first day of adulthood as a consequence of modified neuronal functions and progresses further as a consequence of the stochastic senescence of muscle cells (7). Hundreds of morphological and behavioural features extracted at once from high resolution videos of the worms was previously shown to produce meaningful classes of mutants ^11–13^. We opted for this approach to build an unbiased multiple parameters database precisely describing the phenotypic progression of worms during ageing. We anticipated that this unique database would comprehend the differences and similarities between animals and between physiological and pathological models over their entire lifespan.

The intimate relationship between ageing and neurodegenerative disorders raised the possibility of shared mechanisms and led to the concept of accelerated ageing ^14^. Delineating the phenotypic ageing trajectories would reveal the relationship between pathological and physiological ageing and would challenge the concept of accelerated ageing. Ageing on the same phenotypic trajectory at different speeds would suggest that shared mechanisms affect the worm physiology at different rates. On the contrary, ageing on different trajectories would suggest that distinct mechanisms affect worm physiology in each context. Several *C. elegans* models exist for pathological ageing, each mimicking a muscular or neurodegenerative human disorder by the overexpression of human aggregation-prone proteins in *C. elegans* ^15,16^. As behavioural changes are amongst the first consequences of ageing in *C. elegans* ^6^, we developed a model of neuronal proteinopathy as pathological model. Alpha-Synuclein is an aggregation-prone protein involved in the pathology of Parkinson’s disease. The expression of human alpha-Synuclein mimics pathological aspects of age-related neurodegenerative diseases in all model organisms tested including yeast ^17^, *Drosophila* ^18,19^ and *C. elegans* ^20–22^. Therefore we based our model on the expression of human alpha-Synuclein fused to YFP in the 8 dopaminergic neurons of *C. elegans*. With a multi-parametric analysis of behaviour, we showed that rather than a mere acceleration of the ageing process, disease-model worms follow a divergent trajectory of ageing. We also showed that the pathological state of animals is predictable from day 1 of adulthood and that their health can be quantified through a health index assessing their range of abilities. We conclude that multi-parametric analysis of behaviour is best suited for the analysis of qualitative and quantitative properties of ageing.

## Results and Discussion

### A comprehensive phenotyping for ageing worms

Describing the dynamic changes occurring during the ageing process from a few chosen phenotypes presents a risk of misleading conclusions because of intrinsic bias in the feature selection process. Using a large set of parameter would mitigate this selection bias and take into account compensation effects between parameters. To this end, we used video tracking of single worms and a quantitative phenotyping method able to detect and quantify the subtle and progressive morphological and behavioural changes occurring at the organismal scale during ageing. Per genotype, 20 to 30 individual worms were video-tracked. Each individual was transferred to a fresh plate, habituated to the new plate for 30 minutes before a 15 minutes tracking period. This procedure was repeated every day from the L4 stage to the death of the individual (Fig. 1). Importantly, it implies that the animals experienced a constant high food and low waste environment over their lifespan. The videos were segmented and skeletonized in order to extract a set of 171 morphological, postural and locomotory features ^11^ (Supplementary Table 1). These features comprise the mean, 5^th^ percentile and 95^th^ percentile of 57 core features that capture changes in ageing animals. This unique dataset was used to capture the physiological and pathological ageing of *C. elegans.*

**Figure 1.**
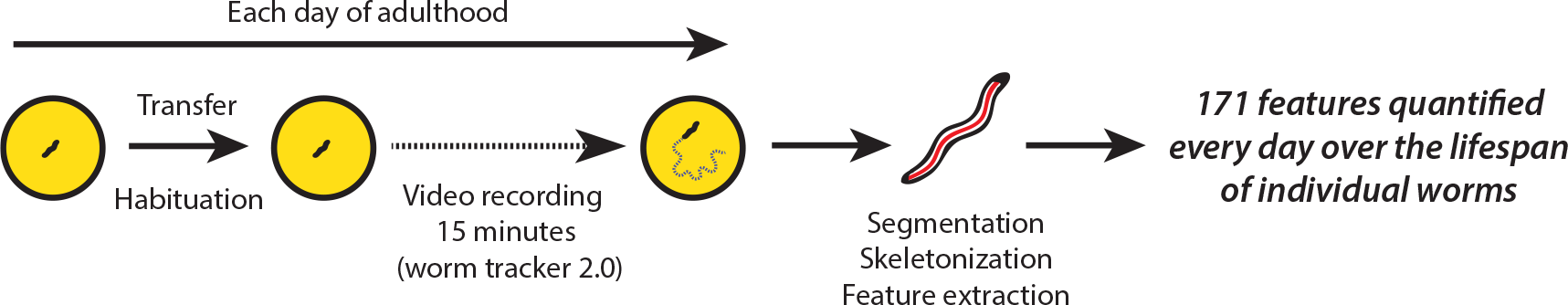
Experimental procedure. Each worm was transferred every day to a fresh plate before habituation and tracking with worm trackers 2.0. Videos were analysed with the worm tracker 2.0 software (segmentation, skeletonization and feature extraction) to extract a set of 171 features.

### Expression of alpha-Synuclein-YFP in the dopaminergic neurons induces pathological ageing

To compare physiological to pathological ageing, we generated a neuron-specific synucleinopathy-like disorder in *C. elegans*. To compensate for possible biases induced by the transgenesis method, we generated several independent integrated transgenic lines expressing YFP (OW953 and OW956) or alpha-Synuclein-YFP (OW939, OW940 and OW949) under the control of the *dat-1* promoter. The *dat-1* promoter is active selectively in the dopaminergic neurons from their specification to the death of the animals. These integrated strains were outcrossed 8 times in the N2 background. In all three strains expressing alpha-Synuclein-YFP, we observed that alpha-Synuclein-YFP progressively relocates into fluorescent foci in the cytoplasm and neurites of the dopaminergic neurons, while YFP remains soluble in control strains (Supplementary Fig. 1a and 1b). Expression of alpha-Synuclein-YFP fusion did not induce neuronal loss of the CEP or ADE dopaminergic neurons until day 20 of adulthood (Supplementary Fig. 1c). One of the 3 strains expressing alpha-Synuclein-YFP weakly but statistically shortened lifespan compared to the wild-type animals while the other 2 strains showed not statistical differences (Supplementary Fig. 1d). Overall, the expression of alpha-Synuclein-YFP in the dopaminergic neurons did not induce other phenotypes quantifiable by hand at the organismal level.

We video-tracked and analysed in detail the phenotype of 3 strains expressing alpha-Synuclein-YFP, 2 strains expressing YFP, the wild-type N2 strain and *cat-2(e1112) II* mutant, a mutant of a tyrosine hydroxylase with decreased levels of dopamine ^23^. The mean value for each of the 171 features was extracted for each strain, for young (L4 stage) and old worms (day 15 of adulthood). Hierarchical clustering was performed using Euclidean distance and between-group average linkage for the entire set of 171 normalized mean values. This unbiased clustering analysis demonstrates a clear phenotypic difference between the L4 stage and the 15 days old worms, independently of their genotype (Fig. 2).

**Figure 2.**
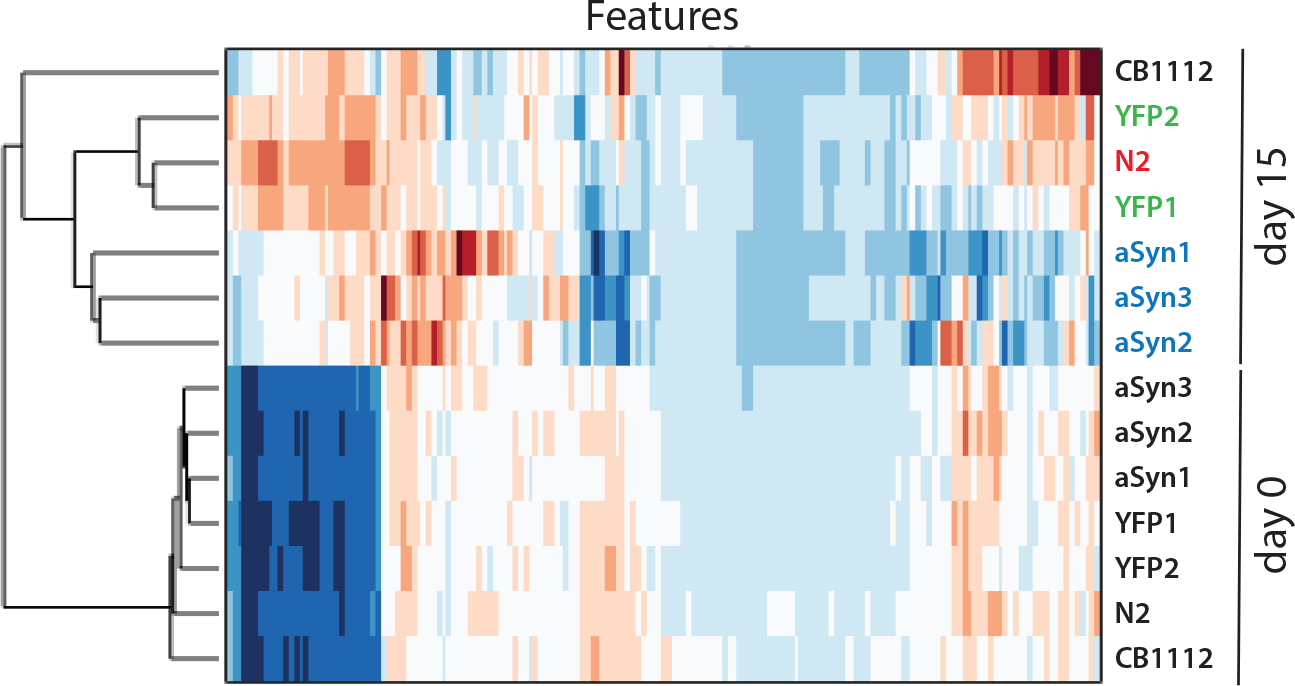
Disease-model and healthy animals are phenotypically different. Hierarchical clustering of the different strains at the L4 stage and day 15 of adulthood. The dataset for 171 parameters and 7 strains was clustered using Euclidean distances and between-group average linkage. For each feature, red and blue shadings indicate means above or below the mean across strains, respectively. The magnitude of the difference is indicated by the shading intensity. Each row represents one strain tracked at L4 (day 0) or day 15 of adulthood. The row labels YFP1, YFP2, aSyn1, aSyn2, aSyn3 are OW953, OW956, OW939, OW940 and OW949 strains, respectively.

Interestingly, while all strains appeared phenotypically similar at the L4 stage, they diverged at day 15. At day 15, the strains expressing YFP clustered with the wild-type animals (N2) whereas the 3 strains expressing alpha-Synuclein-YFP clustered together and showed strong phenotypic differences with YFP and N2 strains. The alpha-Synuclein-YFP-expressing strains also induced a phenotypic signature unrelated to the one observed in the *cat-2* mutant. Therefore, a specific phenotypic signature is induced at day 15 by the expression of the aggregation-prone protein alpha-Synuclein-YFP in dopaminergic neurons. This phenotype is not caused by the mere overexpression of the exogenous fluorophore YFP and appears different to the phenotypes induced by reduced dopamine levels in the *cat-2* mutants. Considering the clustering observed among genotypes at day 15, strains of the same genotype were pooled for subsequent analysis.

### Divergent physiological and pathological ageing trajectories

To select the most relevant features among the 171 for further analysis, those showing the most significant differences between alpha-Synuclein-YFP and YFP strains after ANOVA testing and Bonferroni correction were retained. After feature selection, a set of 47 significant morphological and behavioural features with the lowest p values was kept for analysis of health over age (Supplementary Table 1). To determine when alpha-Synuclein-YFP expression in dopaminergic neurons initiates pathological ageing, the Euclidean distances between each genotype and the N2 controls were calculated for each day of life for the 47 selected features (Fig. 3). Phenotypic differences between alpha-Synuclein-YFP strains and the N2 and YFP controls became significant from day 9 of adulthood and continue to increase with age. This result is consistent with neurodegenerative disorders, in which phenotypes occur with a late onset and worsen over age.

**Figure 3.**
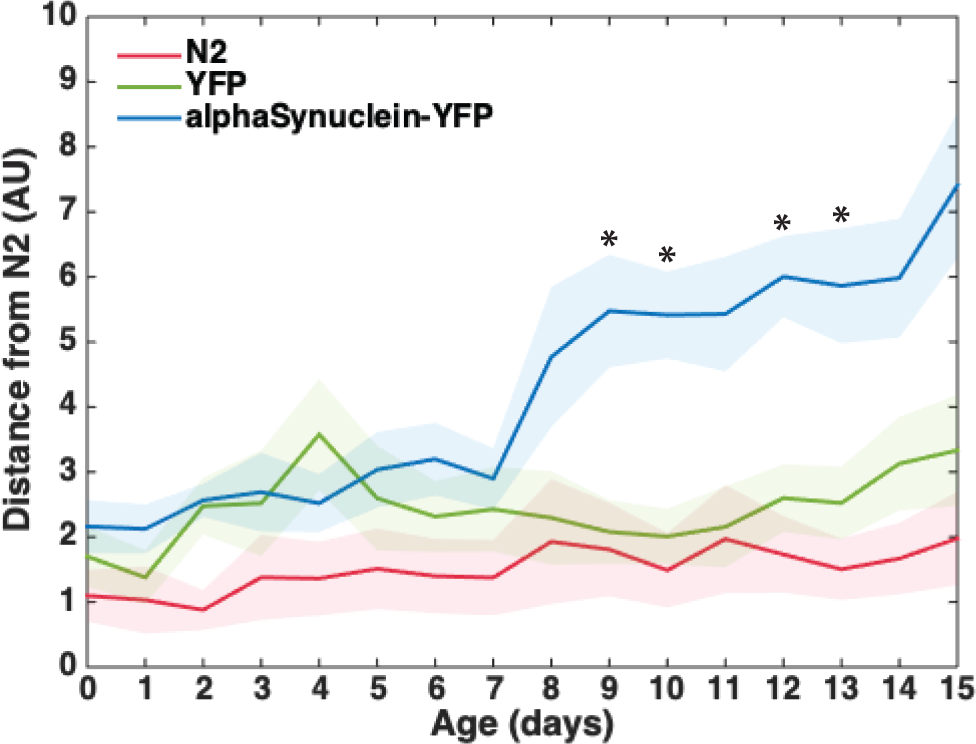
Distance between disease-model and healthy animals increases with age. For the 47 most significant features, the Euclidean distance between each group of strains and N2 was calculated for each day of life by bootstrapping (10 000 random resamplings with replacement) and represented over age. Shaded areas indicate standard deviations. * Indicates the days when the 2 groups YFP and alpha-Synuclein-YFP become significantly different (p value < 0.05, comparison of confidence intervals).

This observation could be the consequence of the acceleration of the physiological ageing process or of a new and divergent trajectory of ageing (Fig. 4a). To explore the later hypothesis, we compared trajectories of ageing of the different genotypes in a phenotypic landscape defined by our 47 parameters. t-SNE (t-distributed Stochastic Neighbour Embedding) was used to visualize the trajectories of ageing in a two-dimensional phenotypic landscape (Fig. 4b). The average behaviour for each genotype at each age was calculated to compare the ageing courses of each genotype. In this 2-dimensional representation of phenotypic ageing, the data appears distributed according to age. It shows a clear evolution of the phenotypes during ageing of wild-type animals, with a trajectory monotonically ranging from young to old animals. The strains expressing YFP follow a trajectory relatively similar to the N2 animals, at the same ageing rate. Instead, the strain expressing alpha-Synuclein-YFP show a clear qualitative shift in their phenotypic profile compared to the controls. In addition, the strains expressing alpha-Synuclein-YFP travelled faster than the N2 strain within the phenotypic landscape. They cluster with the phenotype of old worms from day 9 of adulthood and remain phenotypically similar until their death. Therefore alpha-Synuclein-YFP induces an accelerated ageing on a divergent phenotypic trajectory of ageing (Fig. 4a, box 3).

**Figure 4.**
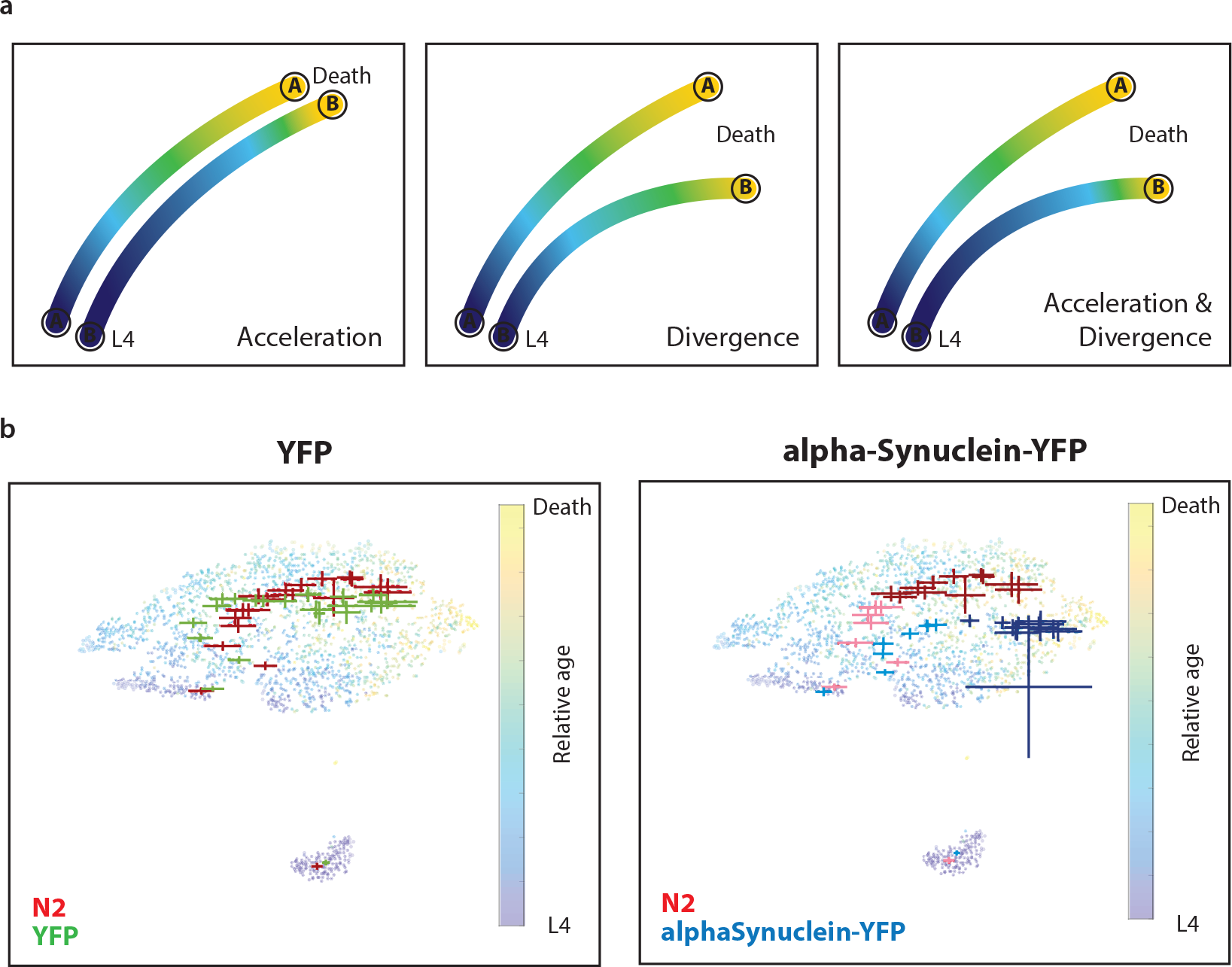
Ageing trajectories differ qualitatively. (a) Examples of comparison of 2 t-SNE trajectories in a 2-D virtual phenotypic space. Colors indicate the relative age of the animals. In the model of accelerated ageing, the 2 trajectories are qualitatively similar but the trajectory B marks an acceleration of the ageing process. In the model of divergent ageing trajectories, the 2 trajectories are qualitatively divergent but the ageing speeds are similar. In the third model, the trajectories are divergents and the trajectory B also shows an acceleration of its ageing process. (b) t-SNE visualization of the behavior of each genotype. Each dot summarizes the phenotype of a single worm at a single age. Colors of the dots indicate the relative age of each single worm, from dark blue (L4) to yellow (death). Trajectories are drawn from the mean and standard error of the mean for each strain at each chronological age. Light-coloured crosses indicate the 8 first days of adulthood while dark-coloured crosses indicate the remaining days of life.

### Multivariate phenotyping predicts the pathological state

To confirm that differences in ageing trajectories are detectable at the phenotypic scale, we classified the genotypes of each animal at each age using the phenotypic data with a binary support vector machine classifier (linear Kernel). From their phenotypes, the genotype of each animal (N2 versus alpha-Synuclein-YFP) is predicted with a high accuracy, with a percentage of good prediction of 86 % across all ages. Importantly, the genotype is predictable from the phenotype of young animals, with 87 % and 90 % of good predictions at day 1 and day 2 of adulthood, respectively. This result highlights the power of multivariate phenotyping for the early detection of pathological states.

### Multivariate recording of phenotypes reveals new ageing markers

As many new parameters were used to describe ageing properties, we also asked whether these features would be new biomarkers of ageing. In fact, 67% of the 171 parameters used in this study highly correlated with age and relative age in non-pathological condition (p value < 0,05 after Bonferroni correction, Pearson’s correlation), and therefore become new biomarkers of *C. elegans* wild-type ageing (Supplementary Table 2). As biomarker of longevity, maximum velocity was previously shown to highly correlate with lifespan ^4^. Interestingly, biomarkers of ageing showing the best correlation coefficients are mainly morphological and postural features, and correlate better with relative age than our approximate value for maximum velocity of the animals.

### Worms expressing alpha-Synuclein lose their behavioural flexibility early and age in an altered health state

As pathological ageing follows a different phenotypic trajectory, defining the “health state” of an individual is not as trivial: any health index must apply to physiological and pathological trajectories of ageing. As any phenotypic parameter is at risk of being characteristic of a specific trajectory, a significant subset of phenotypic parameter is better suited to generate an appropriate index. The distribution of values for each of the 47 parameters over 15 minutes recording covers the dynamic range available to the individual at each age across genotypes. Frailty and ageing could reduce phenotypic complexity as well as the ability of the individuals to produce the strongest - and potentially most fitted - responses to internal or external variations. To assess whether the pathological trajectory correlates with reduced fitness, we defined a health index based on the range of abilities of each animal. This index is calculated as the mean difference of maximum (95^th^ percentile) and minimum (5^th^ percentile) values for all 57 core features. It has the advantage of taking into account the maximum and minimum abilities of each worm for each phenotypic parameter, therefore giving an indication on the global phenotypic abilities of the individuals.

Quantified by the *range of abilities index*, the phenotypic flexibility of the wild-type animals and control strains expressing YFP decreases at low rate over the chronological and relative age (Fig. 5a and 5b). However phenotypic flexibility declines earlier and faster for strains expressing alpha-Synuclein-YFP, diverging from YFP control strains from day 4 of adulthood and worsening at a higher rate from day 5 of adulthood. Using relative age (age relative to lifespan) as a time scale confirms the relationship between flexibility decline and age and the earlier decline for strains expressing alpha-Synuclein-YFP. This result suggests that the phenotype of disease-model worms becomes less flexible than control worms with an early onset.

**Figure 5.**
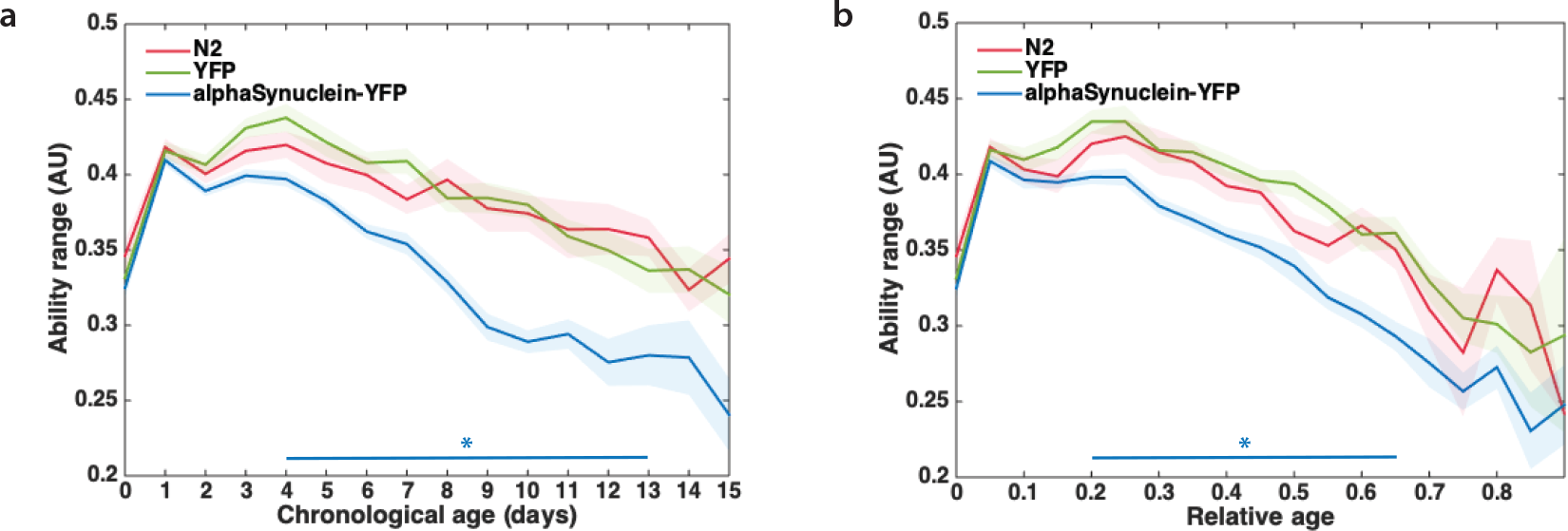
Disease-model worms loose flexibility over age. The ability range was calculated as the mean difference between maximum and minimum values for all features. (a) Evolution of the range of ability over chronological age. (b) Evolution of the range of ability over relative age. * Indicates the days when the 2 groups YFP and alpha-Synuclein-YFP become significantly different (p value < 0.05, multiple t-test with Sidak correction).

### Uncoupling of healthspan and lifespan in a pathological context

In terms of kinetics, physiological and pathological trajectories of ageing correspond to long and progressive decline in health as illustrated by the phenotypic flexibility. Therefore defining cut-offs such as healthspan or gerospan appears artificial. However, at the population level, while having lifespans close to control worms (Supplementary Fig. 1d), the disease model shows an earlier onset of decline in health and spends a longer time unhealthy, supporting the uncoupling of healthspan and lifespan in a pathological context.

### Precise representation of ageing by multivariate approach

Our results underscore the importance of multivariate approaches to take into account the possibility of different ageing trajectories for different genotypes. In *C. elegans*, the previous method based on the aggregation of 5 predictive biomarkers of biological age recorded in one strain did not assess this possibility ^5^. In human, data medicine has defined molecular markers predictive of the biological age such as gene expression ^24^, DNA methylation profiles ^25^ or functional parameters such as physical activity ^26^. Based on one or few of these predictive markers it is possible to quantify the difference between the biological and the chronological age of an individual. This difference is referred as “age acceleration” and is predictive of health risks ^27^. Our results suggest that pathological states do not correspond to such a simple “acceleration of ageing” but rather to a modification of the aging trajectory in a multi-parametric space.

Divergent trajectories of ageing have several implications. First, it suggests that distinct biological mechanisms affect the worm physiology during physiological and pathological ageing. Second, divergent trajectories affect the definition of a generalized health index, as this index would have to apply to all potential trajectories. Using the worm prognosis as it was suggested in (5) might not be the best informative indicator for health. While the alpha-Synuclein-YFP strains have survival curves close to wild-type animals (Supplementary Fig. 1d), our index show consistently worse health for the disease-model worms compared to N2. Finally, we envisage that each age-related disorder might generate a specific ageing trajectory, offering a specific signature for each age-related disorder.

Therefore, our multi-parametric framework offers a better practice to compare genotypic, environmental or inter-individual effects on ageing. We demonstrate that this approach is highly predictive for specific health risks. Coupled to pharmacological or genetic screen, our framework would provide precise read-outs to describe how genetic mutations or therapeutic drugs reshapes ageing trajectories. Altogether, this work highlights the potential of non-invasive recording of multiple health parameters to predict and monitor human trajectories of ageing. Linked to medical records, such collection of multivariate data could be used for the development of personalized medical treatments to prevent or treat age-related pathologies.

## Material and Methods

### Plasmids strains and media

P(dat-1)::YFP and P(dat-1)::alpha-Synuclein-YFP were generated by cloning the P(dat-1) promoter in the plasmids pRP2386 ^28^ and pPD30.38 (Addgene plasmid # 1443) respectively, on place of the P(unc-54) promoter.

The following strains were used: N2 (*wt*), CB1112 (*cat-2(e1112) II*), OW939 (zgIs113[*Pdat-1::alpha-Synuclein::YFP*]), OW940 (zgIs128[*Pdat-1::alpha-Synuclein::YFP*]), OW949 (zgIs125[*Pdat-1::alpha-Synuclein::YFP*]), OW953 (zgIs138[*Pdat-1::YFP*]), OW956 (zgIs144[*Pdat-1::YFP*]), OQ99 (ulbEx[*Pdat-1::mKate*]), AQ2028 (Ex[*Pdat-1::ChR2*;*Punc-122::GFP*]).

Transgenic *C. elegans* (OW939, OW940, OW49, OW953 and OW956) were generated by microinjection and irradiation with gamma rays for integration of the transgene in the genome. Strains were outcrossed 8 times with the N2 strain. Standard conditions were used to maintain and propagate *C. elegans* strains at 20 °C.

### Microscopy

Worms were age-synchronized with egg-laying windows of 4 hours and grown on NGM without FuDR. Worms were transferred on fresh plates everyday until day 3 of adulthood to avoid progeny proliferation. Images were acquired with a Zeiss LSM 780 confocal microscope.

### Counting of dopaminergic neurons

To assess neuronal loss, the strains OW940 and OW953 were crossed with the strain OQ99. Worms were grown on NGM supplemented with FuDR from day 1 until day 20 of adulthood to prevent progeny proliferation. At day 1, 5, 10, 15 and 20 of adulthood, 45 worms were collected and imaged with a Zeiss Axio Zoom V16. CEP and ADE neurons expressing mKate were counted from the acquired pictured.

### Collection of behavioural data

For each strain, between 20 and 30 single worms were video-tracked using previously described Worm Trackers 2.0 ^11^. Worms were tracked longitudinally every day from the L4 stage to death. To avoid environmental variations to bias measurements, genotypes and ages were randomized across 4 trackers and over 3 months of experimentation. Worms were maintained in strict conditions at 20 °C until and during the point of tracking.

Single-worm tracking was performed as previously described ^11,29^. Briefly, 3 cm plates containing low peptone NGM were seeded with 20 μL of OP50 30 minutes prior tracking. Each day, each single worm was picked with a sterile eyelash on a new fleshly seeded plate and let habituate for 30 minutes. Worm behaviour was then recorded for 15 minutes at 25 frames per second. Videos were analysed with the freely available Worm Tracker 2.0 software to extract behavioural features as time series data ^11^. The worm-behaviour data is available on an open-source platform ^13^ (http://movement.openworm.org/).

### Feature extraction and data preparation

Features were extracted from the time series data produced by the Worm Tracker 2.0 software. For each worm, a set of 171 features were extracted as the mean, 5^th^ percentile and 95^th^ percentile of 57 core features of the time-series data. These features contain information about the worm morphology, posture and locomotion.

After feature extraction, a selection of representative features was operated to eliminate non-informative features. ANOVA was performed between two groups of worms, YFP and alpha-Synuclein-YFP, to identify features that show the most significant variation between the two groups. The 47 features with the lowest p value (p value < 10^−8^) were selected to compose the dataset. The dataset was then standardized with z-score to compensate for the different units of each feature.

### Genotype prediction

Genotype prediction was done using a binary support vector machine classifier (linear Kernel). Independently of age or genotype, half of the dataset was randomly selected to train the model and the other half was used for prediction. Prediction was done without cross-validation on the train set.

### Health index

Health index was calculated from the difference of raw maximum (95^th^ percentile) and minimum (5^th^ percentile) values for all 57 core features. The mean difference was computed after normalization (scaling in the range [0,1]) to obtain the health index.

### Lifespan analysis

Lifespan was assessed from the tracking experiment at 20 °C without FuDR. The number of live animals was scored every day until death. Lifespan was analysed by GraphPad Prism analysis software.

## Acknowledgments

pPD30_38 was a gift from Andrew Fire (Addgene plasmid # 1443). We thank Stijn Mouton for reading the manuscript. C.N.M. was the beneficiary of fellowships and travel grants from the Fondation pour la Recherche Médicale (FRM), the Stichting Parkinson Fonds, the Université Libre de Bruxelles (ULB), the International Brain Research Organization (IBRO) and the European Cooperation for Science and Technology (COST-Genie). This project was funded by a European Research Council (ERC) starting grant (to E.A.A.N.), a Meervoud grant from the Division for Earth and Life Sciences (ALW) with financial aid from The Netherlands Organization for Scientific Research (NWO) (to E.A.A.N), the Alumni chapter Gooische Groningers facilitated by the Ubbo Emmius Fonds (to E.A.A.N.).

## Author Contributions

Conceptualization, C.N.M., A.E.X.B., E.A.A.N. and P.L.; Methodology, C.N.M.; Formal Analysis and Investigation, C.N.M., B.B., A.E.X.B. and P.L.; Resources, E.A.A.N., W.R.S. and A.E.X.B.; Writing – Original Draft, C.N.M.; Writing – Review and Editing, C.N.M, P.L., A.E.X.B, E.A.A.N and W.R.S.; Supervision, E.A.A.N., A.E.X.B and P.L; Funding Acquisition, C.N.M, E.A.A.N. and P.L.

## References

1. Kenyon, C. The Plasticity of Aging: Insights from Long-Lived Mutants. Cell 120, 449–460 (2005).

2. Kenyon, C. J. The genetics of ageing. Nature 464, 504–512 (2010).

3. Bansal, A., Zhu, L. J., Yen, K. & Tissenbaum, H. A. Uncoupling lifespan and healthspan in Caenorhabditis elegans longevity mutants. Proc. Natl. Acad. Sci. 112, E277–E286 (2015).

4. Hahm, J.-H. et al. *C. elegans* maximum velocity correlates with healthspan and is maintained in worms with an insulin receptor mutation. Nat. Commun. 6, 8919 (2015).

5. Zhang, W. B. et al. Extended Twilight among Isogenic C. elegans Causes a Disproportionate Scaling between Lifespan and Health. Cell Syst. 3, 333–345.e4 (2016).

6. Croll, N. A., Smith, J. M. & Zuckerman, B. M. The aging process of the nematode Caenorhabditis elegans in bacterial and axenic culture. Exp. Aging Res. 3, 175–189 (1977).

7. Herndon, L. A. et al. Stochastic and genetic factors influence tissue-specific decline in ageing *C. elegans*. Nature 419, 808–814 (2002).

8. Garigan, D. et al. Genetic Analysis of Tissue Aging in Caenorhabditis elegans: A Role for Heat-Shock Factor and Bacterial Proliferation. Genetics 161, 1101–1112 (2002).

9. McGee, M. D. et al. Loss of intestinal nuclei and intestinal integrity in aging C. elegans. Aging Cell 10, 699–710 (2011).

10. Toth, M. L. et al. Neurite Sprouting and Synapse Deterioration in the Aging Caenorhabditis elegans Nervous System. J. Neurosci. 32, 8778–8790 (2012).

11. Yemini, E., Jucikas, T., Grundy, L. J., Brown, A. E. X. & Schafer, W. R. A database of *Caenorhabditis elegans* behavioral phenotypes. Nat. Methods 10, 877–879 (2013).

12. Javer Avelino, Ripoll-Sánchez Lidia & Brown André E.X. Powerful and interpretable behavioural features for quantitative phenotyping of Caenorhabditis elegans. Philos. Trans. R. Soc. B Biol. Sci. 373, 20170375 (2018).

13. Javer, A. et al. An open-source platform for analyzing and sharing worm-behavior data. Nat. Methods 15, 645–646 (2018).

14. Kennedy, B. K. et al. Geroscience: Linking Aging to Chronic Disease. Cell 159, 709–713 (2014).

15. Sin, O., Michels, H. & Nollen, E. A. A. Genetic screens in Caenorhabditis elegans models for neurodegenerative diseases. Biochim. Biophys. Acta BBA - Mol. Basis Dis. 1842, 1951–1959 (2014).

16. Cooper, J. F. & Van Raamsdonk, J. M. Modeling Parkinson’s Disease in C. elegans. J. Park. Dis. 8, 17–32 (2018).

17. Willingham, S., Outeiro, T. F., DeVit, M. J., Lindquist, S. L. & Muchowski, P. J. Yeast genes that enhance the toxicity of a mutant huntingtin fragment or alpha-synuclein. Science 302, 1769–1772 (2003).

18. Feany, M. B. & Bender, W. W. A Drosophila model of Parkinson’s disease. Nature 404, 394–398 (2000).

19. Dabool, L., Juravlev, L., Hakim-Mishnaevski, K. & Kurant, E. Modeling Parkinson’s disease in adult Drosophila. J. Neurosci. Methods 311, 89–94 (2019).

20. Lakso, M. et al. Dopaminergic neuronal loss and motor deficits in Caenorhabditis elegans overexpressing human α-synuclein. J. Neurochem. 86, 165–172 (2003).

21. Cao, S., Gelwix, C. C., Caldwell, K. A. & Caldwell, G. A. Torsin-Mediated Protection from Cellular Stress in the Dopaminergic Neurons of Caenorhabditis elegans. J. Neurosci. 25, 3801–3812 (2005).

22. Kuwahara, T. et al. Familial Parkinson Mutant α-Synuclein Causes Dopamine Neuron Dysfunction in Transgenic Caenorhabditis elegans. J. Biol. Chem. 281, 334–340 (2006).

23. Sanyal, S. et al. Dopamine modulates the plasticity of mechanosensory responses in Caenorhabditis elegans. EMBO J. 23, 473–482 (2004).

24. Peters, M. J. et al. The transcriptional landscape of age in human peripheral blood. Nat. Commun. 6, 8570 (2015).

25. Hannum, G. et al. Genome-wide Methylation Profiles Reveal Quantitative Views of Human Aging Rates. Mol. Cell 49, 359–367 (2013).

26. Pyrkov, T. V. et al. Quantitative characterization of biological age and frailty based on locomotor activity records. Aging 10, 2973–2990 (2018).

27. Dugué, P.-A. et al. Association of DNA Methylation-Based Biological Age With Health Risk Factors and Overall and Cause-Specific Mortality. Am. J. Epidemiol. 187, 529–538 (2018).

28. van Ham, T. J. et al. C. elegans Model Identifies Genetic Modifiers of α-Synuclein Inclusion Formation During Aging. PLOS Genet. 4, e1000027 (2008).

29. Yemini, E., Kerr, R. A. & Schafer, W. R. Preparation of Samples for Single-Worm Tracking. Cold Spring Harb. Protoc. 2011, 1475–1479 (2011).

