## Supplementary Information for "Deep behavioural phenotyping reveals divergent trajectories of ageing and quantifies health state in *C. elegans*"

#### Supplementary Figure 1

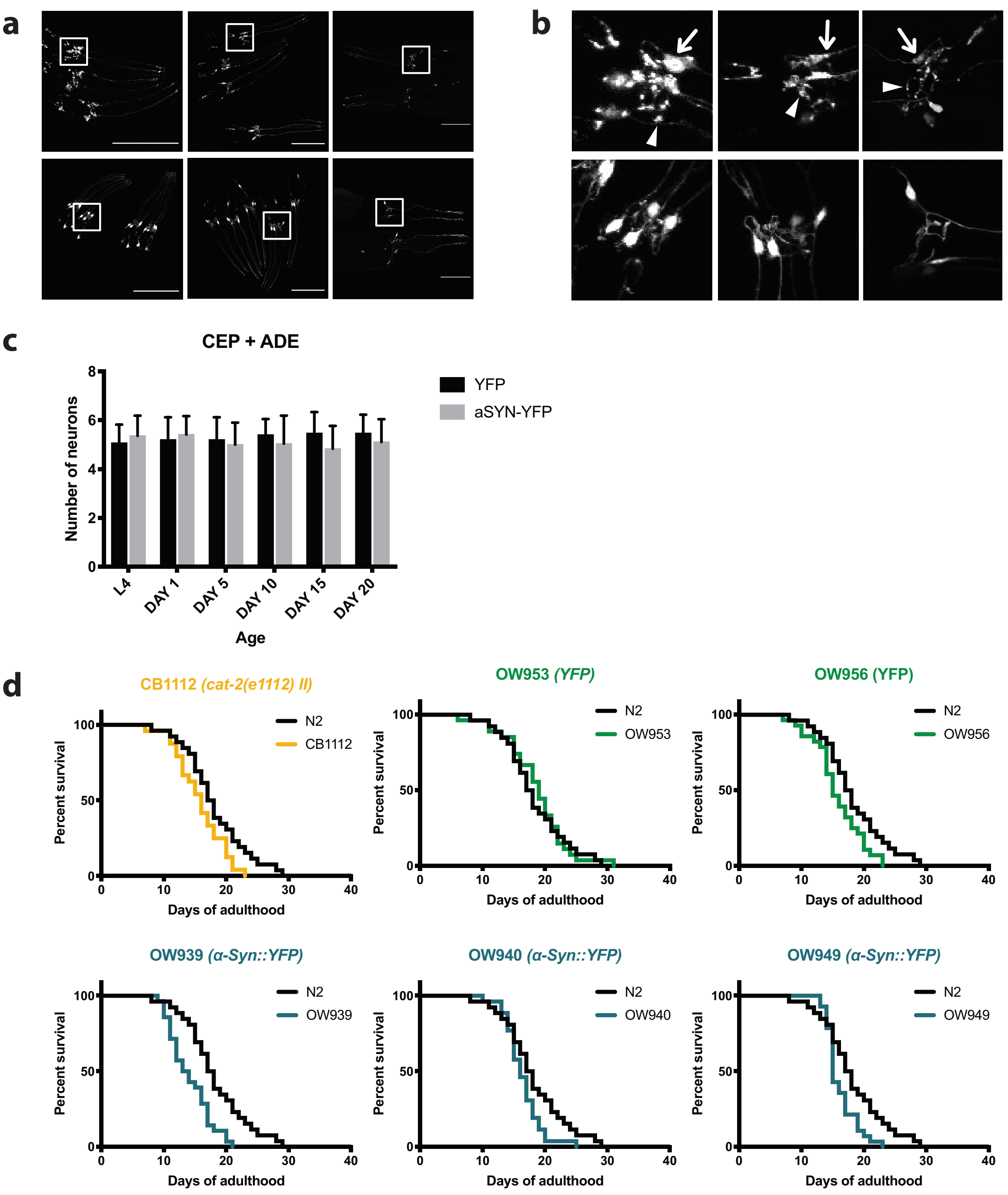


| Strain | N2 | CB1112 | OW953 | OW956 | OW939 | OW940 | OW949 |
|---|---|---|---|---|---|---|---|
| Genotype | *wt* | *cat-2(e1112) II* | *$P_{dat-1}$::YFP* | *$P_{dat-1}$::YFP* | *$P_{dat-1}$::α-Syn::YFP* | *$P_{dat-1}$::α-Syn::YFP* | *$P_{dat-1}$::α-Syn::YFP* |
| Number of death events | 26 | 24 | 27 | 28 | 28 | 26 | 28 |
| Median survival (days) | 17,5 | 16 | 19 | 15 | 13,5 | 16 | 15 |
| Curve comparison to N2 (Wilcoxon) | na | ns | ns | ns | ** (p < 0,005) | ns | ns |

**Supplementart Figure 1. Characterisation of the synucleinopathy model.** (a) Confocal laser scanning images showing the expression of YFP and of alpha-Synuclein-YFP in the head region of transgenic animals at different ages. Representative pictures for each genotype. aSyn: alpha-Synuclein-YFP. (b) Magnification of the corresponding inserts of (a). Arrows and arrowheads indicate protein inclusions in somas and neurites, respectively. (c) Count of CEP and ADE neurons over age do not show neuronal loss. n = 45 animals per age. Bars indicate standard deviations. YFP and aSYN-YFP are the OW953 and OW940 strains, respectively. (d) Survival curves of N2, *cat-2(e1112)II*, YFP-expressing and alpha-Synuclein-expressing worms. Results from the longitudinal study of behaviour. Statistical results from curves comparisons to N2.

#### Supplementary Table 1

**Phenotypic features used in this study.** Results of ANOVA testing for all features. Corrected p values after Bonferroni correction are shown. The top 47 features (p value < $10^{-8}$) selected for strains comparisons are highlighted in grey.

| Features names | p Values | Features names | p Values |
|---|---|---|---|
| **'worm.posture.amplitude.max_Q95'** | **1,74E-26** | 'worm.morphology.width.midbody_Mean' | 0,0012692 |
| **'worm.morphology.length_Q95'** | **3,76E-26** | 'worm.morphology.width.head_Mean' | 0,0014140 |
| **'worm.morphology.length_Mean'** | **4,21E-25** | 'absolute worm.locomotion.velocity.tailTip.speed_Q05' | 0,0014976 |
| **'worm.morphology.length_Q05'** | **4,24E-23** | 'worm.posture.kinks_Q05' | 0,0017034 |
| **'worm.posture.tracklength_Q95'** | **8,21E-23** | 'backward worm.locomotion.velocity.midbody.speed_Q05' | 0,0019002 |
| **'worm.morphology.area_Q95'** | **1,18E-22** | 'absolute worm.posture.bends.hips.mean_Mean' | 0,0019523 |
| **'forward worm.locomotion.velocity.headTip.speed_Q95'** | **4,99E-22** | 'absolute worm.posture.bends.neck.mean_Mean' | 0,0020771 |
| **'absolute worm.locomotion.velocity.headTip.speed_Q95'** | **5,13E-21** | 'absolute worm.path.curvature_Q05' | 0,0038361 |
| **'worm.posture.wavelength.secondary_Q95'** | **2,60E-20** | 'worm.morphology.width.tail_Mean' | 0,0055987 |
| **'worm.posture.amplitude.max_Mean'** | **2,77E-20** | 'worm.morphology.width.head_Q05' | 0,0064034 |
| **'forward worm.locomotion.velocity.head.speed_Q95'** | **5,46E-20** | 'absolute worm.locomotion.bends.tail.amplitude_Mean' | 0,0065738 |
| **'worm.posture.wavelength.secondary_Mean'** | **2,43E-19** | 'worm.posture.eccentricity_Mean' | 0,0078757 |
| **'absolute worm.locomotion.velocity.head.speed_Q95'** | **6,85E-19** | 'worm.path.range_Mean' | 0,0079353 |
| **'worm.morphology.area_Mean'** | **9,09E-19** | 'absolute worm.locomotion.bends.foraging.angleSpeed_Q05' | 0,0092042 |
| **'backward worm.locomotion.velocity.headTip.speed_Q95'** | **5,22E-18** | 'absolute worm.posture.bends.head.stdDev_Q95' | 0,0140647 |
| **'worm.posture.wavelength.secondary_Q05'** | **3,48E-16** | 'absolute worm.locomotion.velocity.head.speed_Q05' | 0,0141741 |
| **'worm.morphology.area_Q05'** | **2,14E-15** | 'absolute worm.posture.bends.head.mean_Q95' | 0,0200830 |
| **'worm.posture.wavelength.primary_Q95'** | **3,16E-15** | 'forward worm.locomotion.velocity.tailTip.speed_Q05' | 0,0244223 |
| **'absolute worm.posture.bends.head.stdDev_Q05'** | **5,13E-15** | 'absolute worm.posture.bends.midbody.mean_Mean' | 0,0291428 |
| **'forward worm.locomotion.velocity.headTip.speed_Mean'** | **6,95E-15** | 'forward worm.locomotion.velocity.head.speed_Q05' | 0,0443128 |
| **'absolute worm.posture.bends.hips.mean_Q95'** | **2,58E-14** | 'absolute worm.locomotion.bends.head.amplitude_Mean' | 0,0568114 |
| **'absolute worm.locomotion.velocity.headTip.speed_Mean'** | **4,68E-14** | 'worm.morphology.areaPerLength_Q05' | 0,0589236 |
| **'worm.posture.wavelength.primary_Mean'** | **6,19E-14** | 'absolute worm.posture.bends.neck.stdDev_Q95' | 0,0953845 |
| **'worm.posture.tracklength_Mean'** | **2,82E-13** | 'worm.morphology.width.midbody_Q05' | 0,1020166 |
| **'forward worm.locomotion.velocity.tailTip.speed_Q95'** | **6,85E-13** | 'absolute worm.posture.bends.tail.stdDev_Mean' | 0,1433990 |
| **'backward worm.locomotion.velocity.headTip.speed_Mean'** | **1,08E-12** | 'absolute worm.posture.bends.midbody.stdDev_Q95' | 0,1700487 |
| **'absolute worm.posture.bends.tail.mean_Q95'** | **1,63E-12** | 'absolute worm.posture.bends.tail.mean_Mean' | 0,3200189 |
| **'forward worm.locomotion.velocity.midbody.speed_Q95'** | **2,97E-12** | 'absolute [illegible] | 0,3422812 |
| **'forward worm.locomotion.velocity.head.speed_Mean'** | **3,40E-12** | 'absolute worm.posture.bends.tail.mean_Q05' | 0,3607064 |
| **'absolute worm.locomotion.velocity.tailTip.speed_Q95'** | **4,93E-12** | 'worm.posture.kinks_Mean' | 0,4632927 |
| **'absolute worm.posture.bends.neck.mean_Q95'** | **6,47E-12** | 'worm.path.range_Q05' | 0,6137305 |
| **'absolute worm.locomotion.velocity.midbody.speed_Q95'** | **8,02E-12** | 'worm.posture.amplitude.ratio_Q95' | 0,9315415 |
| **'absolute worm.locomotion.velocity.head.speed_Mean'** | **1,39E-11** | 'worm.posture.tracklength_Q05' | 1,1460802 |
| **'backward worm.locomotion.velocity.head.speed_Q95'** | **3,57E-11** | 'absolute worm.posture.directions.head_Q95' | 1,3686015 |
| **'absolute worm.posture.bends.neck.stdDev_Q05'** | **3,68E-11** | 'absolute worm.locomotion.bends.foraging.angleSpeed_Q95' | 1,4210454 |
| **'absolute worm.locomotion.velocity.tail.speed_Q95'** | **4,64E-11** | 'absolute worm.posture.directions.head_Q05' | 1,8789012 |
| **'worm.posture.eigenProjection2_Q05'** | **5,91E-10** | 'absolute worm.posture.bends.tail.stdDev_Q95' | 2,0253626 |
| **'backward worm.locomotion.velocity.head.speed_Mean'** | **6,72E-10** | 'worm.morphology.width.tail_Q05' | 2,1484318 |
| **'worm.posture.eigenProjection3_Q05'** | **9,72E-10** | 'forward worm.locomotion.velocity.midbody.speed_Q05' | 2,4939381 |
| **'absolute worm.locomotion.bends.tail.amplitude_Q95'** | **1,33E-09** | 'absolute worm.posture.directions.tail_Q05' | 2,5828003 |
| **'backward worm.locomotion.velocity.tailTip.speed_Q05'** | **2,43E-09** | 'absolute worm.locomotion.velocity.midbody.speed_Q05' | 2,8130685 |
| **'absolute worm.locomotion.bends.midbody.amplitude_Q95'** | **3,04E-09** | 'absolute worm.locomotion.velocity.tail.speed_Q05' | 2,9168676 |
| **'backward worm.locomotion.velocity.midbody.speed_Q95'** | **7,26E-09** | 'absolute worm.posture.bends.midbody.mean_Q05' | 3,5399417 |
| **'worm.posture.wavelength.primary_Q05'** | **1,15E-08** | 'worm.posture.eigenProjection6_Mean' | 3,6767931 |
| **'absolute worm.posture.bends.midbody.mean_Q95'** | **1,26E-08** | 'forward worm.locomotion.velocity.tail.speed_Q05' | 3,9217023 |
| **'forward worm.locomotion.velocity.tailTip.speed_Mean'** | **3,80E-08** | 'worm.posture.eigenProjection4_Q95' | 4,0320272 |
| **'backward worm.locomotion.velocity.tailTip.speed_Q95'** | **4,79E-08** | 'worm.posture.eigenProjection6_Q05' | 4,7321300 |
| 'forward worm.locomotion.velocity.midbody.speed_Mean' | 1,23E-07 | 'absolute worm.posture.directions.tail_Q95' | 5,2551285 |
| 'backward worm.locomotion.velocity.tail.speed_Q95' | 1,43E-07 | 'worm.posture.eccentricity_Q95' | 6,3400845 |
| 'forward worm.locomotion.velocity.tail.speed_Q95' | 1,59E-07 | 'absolute worm.locomotion.bends.tail.frequency_Q05' | 6,7977750 |
| 'absolute worm.locomotion.velocity.tailTip.speed_Mean' | 1,97E-07 | 'absolute worm.posture.bends.head.mean_Q05' | 8,4162572 |
| 'absolute worm.posture.bends.neck.stdDev_Mean' | 1,98E-07 | 'absolute worm.posture.bends.hips.stdDev_Q05' | 10,253462 |
| 'absolute worm.locomotion.bends.midbody.amplitude_Mean' | 2,26E-07 | 'absolute worm.locomotion.bends.midbody.frequency_Q05' | 14,417122 |
| 'worm.morphology.widthPerLength_Q05' | 2,31E-07 | 'worm.posture.amplitude.ratio_Mean' | 14,943425 |
| 'worm.posture.eigenProjection3_Q95' | 3,16E-07 | 'worm.posture.eigenProjection4_Q05' | 15,394893 |
| 'absolute worm.locomotion.velocity.midbody.speed_Mean' | 6,61E-07 | 'absolute worm.locomotion.bends.foraging.amplitude_Q05' | 20,793995 |
| 'worm.morphology.width.midbody_Q95' | 1,08E-06 | 'absolute worm.locomotion.bends.midbody.frequency_Q95' | 28,580525 |
| 'worm.morphology.widthPerLength_Mean' | 1,25E-06 | 'absolute worm.posture.bends.midbody.stdDev_Mean' | 30,839465 |
| 'forward worm.locomotion.velocity.tail.speed_Mean' | 1,61E-06 | 'absolute worm.locomotion.bends.head.frequency_Q05' | 31,005707 |
| 'worm.morphology.areaPerLength_Q95' | 2,32E-06 | 'absolute worm.locomotion.bends.head.frequency_Q95' | 35,305994 |

| | | | |
|---|---|---|---|
| 'backward worm.locomotion.velocity.headTip.speed_Q05' | 2,71E-06 | 'absolute worm.posture.bends.hips.mean_Q05' | 36,229061 |
| 'absolute worm.locomotion.velocity.tail.speed_Mean' | 4,26E-06 | 'absolute worm.locomotion.bends.head.amplitude_Q05' | 36,894643 |
| 'worm.posture.eccentricity_Q05' | 6,54E-06 | 'worm.posture.kinks_Q95' | 45,206222 |
| 'backward worm.locomotion.velocity.tailTip.speed_Mean' | 9,04E-06 | 'worm.posture.eigenProjection5_Q95' | 45,718379 |
| 'worm.posture.eigenProjection2_Q95' | 1,47E-05 | 'worm.posture.eigenProjection2_Mean' | 46,168184 |
| 'absolute worm.path.curvature_Mean' | 2,27E-05 | 'absolute worm.locomotion.bends.tail.frequency_Q95' | 52,813227 |
| 'worm.posture.amplitude.max_Q05' | 2,92E-05 | 'worm.posture.eigenProjection5_Q05' | 54,141731 |
| 'absolute worm.locomotion.velocity.headTip.speed_Q05' | 2,95E-05 | 'worm.posture.eigenProjection3_Mean' | 63,296024 |
| 'absolute worm.posture.bends.head.stdDev_Mean' | 3,64E-05 | 'absolute worm.posture.bends.neck.mean_Q05' | 68,102927 |
| 'absolute worm.locomotion.bends.foraging.amplitude_Mean' | 5,18E-05 | 'absolute worm.posture.bends.hips.stdDev_Mean' | 78,117391 |
| 'forward worm.locomotion.velocity.headTip.speed_Q05' | 5,94E-05 | 'absolute worm.locomotion.bends.midbody.frequency_Mean' | 84,442187 |
| 'worm.morphology.width.head_Q95' | 6,37E-05 | 'absolute worm.locomotion.bends.head.frequency_Mean' | 94,306840 |
| 'worm.morphology.width.tail_Q95' | 8,12E-05 | 'worm.posture.eigenProjection4_Mean' | 99,059179 |
| 'backward worm.locomotion.velocity.head.speed_Q05' | 0,0001017 | 'worm.posture.eigenProjection5_Mean' | 106,44922 |
| 'worm.path.range_Q95' | 0,0001141 | 'absolute worm.locomotion.bends.tail.amplitude_Q05' | 109,67484 |
| 'worm.morphology.widthPerLength_Q95' | 0,0001178 | 'absolute worm.posture.bends.hips.stdDev_Q95' | 110,79533 |
| 'absolute worm.locomotion.bends.midbody.amplitude_Q05' | 0,0001725 | 'absolute worm.posture.bends.midbody.stdDev_Q05' | 121,44503 |
| 'absolute worm.path.curvature_Q95' | 0,0001917 | 'worm.posture.eigenProjection1_Mean' | 132,72398 |
| 'backward worm.locomotion.velocity.tail.speed_Mean' | 0,0002473 | 'absolute worm.locomotion.bends.tail.frequency_Mean' | 137,90170 |
| 'backward worm.locomotion.velocity.midbody.speed_Mean' | 0,0002947 | 'absolute worm.posture.directions.head_Mean' | 140,13290 |
| 'absolute worm.locomotion.bends.head.amplitude_Q95' | 0,0005141 | 'absolute worm.posture.bends.tail.stdDev_Q05' | 147,21234 |
| 'worm.morphology.areaPerLength_Mean' | 0,0007337 | 'absolute worm.posture.directions.tail_Mean' | 147,30753 |
| 'worm.posture.eigenProjection1_Q95' | 0,0008453 | 'absolute worm.posture.bends.head.mean_Mean' | 150,91840 |
| 'backward worm.locomotion.velocity.tail.speed_Q05' | 0,0009876 | 'worm.posture.eigenProjection6_Q95' | 152,24354 |
| 'absolute worm.locomotion.bends.foraging.amplitude_Q95' | 0,0010336 | 'worm.posture.amplitude.ratio_Q05' | 159,95099 |
| 'worm.posture.eigenProjection1_Q05' | 0,0011119 | | |

### Supplementary Table 2

**Correlation of features to age and relative age.** Pearson's correlation coefficient to age and relative age for each of the features used in this study.

| Features names | Corr. to age | Corr. to rel. age | Features names | Corr. to age | Corr. to rel. age |
|---|---|---|---|---|---|
| 'worm.morphology.area_Q05' | 0,7341 | 0,7802 | 'backward worm.locomotion.velocity.head.speed_Q05' | -0,1297 | -0,1949 |
| 'worm.morphology.area_Mean' | 0,7126 | 0,7634 | 'absolute worm.posture.bends.tail.stdDev_Mean' | -0,1430 | -0,1054 |
| 'worm.posture.wavelength.primary_Mean' | 0,7119 | 0,7428 | 'absolute worm.posture.bends.hips.stdDev_Q05' | -0,1434 | -0,1151 |
| 'worm.morphology.areaPerLength_Q05' | 0,6965 | 0,7390 | 'absolute worm.locomotion.bends.midbody.frequency_Q95' | -0,1454 | -0,1578 |
| 'worm.morphology.area_Q95' | 0,6900 | 0,7453 | 'absolute worm.posture.bends.neck.mean_Q05' | -0,1456 | -0,0924 |
| 'worm.morphology.width.midbody_Q05' | 0,6849 | 0,7296 | 'absolute worm.locomotion.velocity.headTip.speed_Q05' | -0,1510 | -0,2173 |
| 'worm.morphology.width.head_Q05' | 0,6629 | 0,7068 | 'absolute worm.locomotion.bends.foraging.angleSpeed_Q05' | -0,1567 | -0,2369 |
| 'worm.posture.wavelength.primary_Q05' | 0,6615 | 0,6959 | 'absolute worm.posture.bends.head.stdDev_Mean' | -0,1571 | -0,1739 |
| 'worm.morphology.areaPerLength_Mean' | 0,6586 | 0,7073 | 'absolute worm.locomotion.bends.midbody.frequency_Q05' | -0,1572 | -0,1129 |
| 'worm.posture.tracklength_Q95' | 0,6407 | 0,6643 | 'worm.posture.kinks_Q05' | -0,1738 | -0,2054 |
| 'worm.morphology.width.head_Mean' | 0,6404 | 0,6757 | 'forward worm.locomotion.velocity.headTip.speed_Q05' | -0,1770 | -0,2389 |
| 'worm.morphology.width.midbody_Mean' | 0,6378 | 0,6923 | 'absolute worm.posture.bends.neck.stdDev_Mean' | -0,1977 | -0,1366 |
| 'worm.morphology.length_Q95' | 0,6377 | 0,6822 | 'absolute worm.locomotion.velocity.head.speed_Q05' | -0,2005 | -0,2612 |
| 'worm.morphology.length_Q05' | 0,6372 | 0,6814 | 'absolute worm.locomotion.velocity.midbody.speed_Q05' | -0,2081 | -0,2620 |
| 'worm.morphology.length_Mean' | 0,6330 | 0,6785 | 'absolute worm.posture.bends.midbody.mean_Mean' | -0,2099 | -0,2057 |
| 'worm.morphology.width.tail_Mean' | 0,6274 | 0,6585 | 'absolute worm.locomotion.velocity.tail.speed_Q05' | -0,2121 | -0,2708 |
| 'worm.morphology.areaPerLength_Q95' | 0,6176 | 0,6717 | 'absolute worm.locomotion.bends.tail.frequency_Mean' | -0,2148 | -0,2388 |
| 'worm.morphology.width.tail_Q05' | 0,6171 | 0,6598 | 'absolute worm.locomotion.bends.head.frequency_Mean' | -0,2161 | -0,2155 |
| 'worm.posture.tracklength_Mean' | 0,6159 | 0,6366 | 'forward worm.locomotion.velocity.head.speed_Q05' | -0,2253 | -0,2815 |
| 'worm.posture.wavelength.secondary_Q95' | 0,6151 | 0,6473 | 'forward worm.locomotion.velocity.tail.speed_Q05' | -0,2267 | -0,2812 |
| 'worm.posture.wavelength.secondary_Mean' | 0,6136 | 0,6390 | 'forward worm.locomotion.velocity.midbody.speed_Q05' | -0,2333 | -0,2840 |
| 'worm.morphology.width.tail_Q95' | 0,5962 | 0,6169 | 'worm.posture.amplitude.ratio_Q95' | -0,2344 | -0,2345 |
| 'worm.morphology.width.head_Q95' | 0,5956 | 0,6251 | 'absolute worm.posture.bends.midbody.mean_Q95' | -0,2391 | -0,2292 |
| 'worm.posture.wavelength.secondary_Q05' | 0,5861 | 0,6068 | 'absolute worm.posture.bends.head.mean_Mean' | -0,2394 | -0,2195 |
| 'worm.morphology.width.midbody_Q95' | 0,5848 | 0,6443 | 'absolute worm.posture.directions.head_Q95' | -0,2410 | -0,2092 |
| 'worm.posture.tracklength_Q05' | 0,5423 | 0,5489 | 'worm.posture.eigenProjection2_Q95' | -0,2421 | -0,2561 |
| 'worm.posture.wavelength.primary_Q95' | 0,5358 | 0,5922 | 'absolute worm.locomotion.velocity.tailTip.speed_Q05' | -0,2426 | -0,2961 |
| 'absolute worm.path.curvature_Mean' | 0,4592 | 0,4758 | 'absolute worm.locomotion.bends.foraging.amplitude_Mean' | -0,2438 | -0,2595 |
| 'absolute worm.path.curvature_Q05' | 0,4407 | 0,4323 | 'worm.posture.eigenProjection3_Q95' | -0,2452 | -0,2504 |
| 'worm.posture.amplitude.max_Mean' | 0,3845 | 0,4295 | 'absolute worm.posture.directions.tail_Q95' | -0,2612 | -0,2718 |
| 'absolute worm.path.curvature_Q95' | 0,3837 | 0,4162 | 'absolute worm.posture.bends.hips.stdDev_Q95' | -0,2640 | -0,2371 |
| 'worm.posture.amplitude.max_Q95' | 0,3714 | 0,3934 | 'absolute worm.posture.bends.neck.stdDev_Q95' | -0,2661 | -0,1839 |
| 'absolute worm.posture.directions.head_Q05' | 0,2970 | 0,2926 | 'absolute worm.posture.bends.neck.mean_Q95' | -0,2665 | -0,2012 |
| 'worm.morphology.widthPerLength_Q05' | 0,2938 | 0,3095 | 'absolute worm.locomotion.bends.foraging.angleSpeed_Mean' | -0,2690 | -0,3424 |
| 'worm.posture.eigenProjection3_Q05' | 0,2810 | 0,2810 | 'absolute worm.locomotion.bends.foraging.angleSpeed_Q95' | -0,2693 | -0,3311 |
| 'absolute worm.posture.directions.tail_Q05' | 0,2568 | 0,2448 | 'forward worm.locomotion.velocity.tailTip.speed_Q05' | -0,2763 | -0,3230 |
| 'worm.posture.amplitude.max_Q05' | 0,2482 | 0,3328 | 'absolute worm.posture.bends.hips.stdDev_Mean' | -0,2764 | -0,2475 |
| 'absolute worm.posture.bends.tail.mean_Q05' | 0,1969 | 0,2731 | 'forward worm.locomotion.velocity.headTip.speed_Mean' | -0,2881 | -0,3216 |
| 'worm.morphology.widthPerLength_Mean' | 0,1728 | 0,1961 | 'absolute worm.locomotion.bends.head.amplitude_Q05' | -0,3023 | -0,2463 |
| 'worm.posture.kinks_Q95' | 0,1642 | 0,1567 | 'absolute worm.locomotion.velocity.headTip.speed_Mean' | -0,3044 | -0,3374 |
| 'worm.posture.eigenProjection4_Q05' | 0,1632 | 0,1617 | 'absolute worm.locomotion.bends.midbody.frequency_Mean' | -0,3187 | -0,2930 |
| 'worm.posture.eigenProjection2_Q05' | 0,1561 | 0,1294 | 'absolute worm.posture.bends.hips.mean_Mean' | -0,3189 | -0,2566 |
| 'absolute worm.locomotion.bends.foraging.amplitude_Q95' | 0,1444 | 0,1334 | 'backward worm.locomotion.velocity.headTip.speed_Mean' | -0,3204 | -0,3510 |
| 'worm.posture.eigenProjection6_Mean' | 0,1296 | 0,1864 | 'backward worm.locomotion.velocity.tailTip.speed_Mean' | -0,3211 | -0,3659 |
| 'absolute worm.posture.bends.midbody.mean_Q05' | 0,1038 | 0,1068 | 'forward worm.locomotion.velocity.headTip.speed_Q95' | -0,3215 | -0,3315 |
| 'worm.posture.eigenProjection6_Q95' | 0,0861 | 0,1534 | 'backward worm.locomotion.velocity.midbody.speed_Mean' | -0,3478 | -0,3941 |
| 'absolute worm.posture.bends.hips.mean_Q05' | 0,0842 | 0,1316 | 'absolute worm.locomotion.bends.head.amplitude_Q95' | -0,3483 | -0,2940 |
| 'absolute worm.posture.bends.tail.mean_Mean' | 0,0815 | 0,1923 | 'absolute worm.locomotion.velocity.headTip.speed_Q95' | -0,3547 | -0,3648 |
| 'worm.posture.eigenProjection6_Q05' | 0,0785 | 0,0857 | 'backward worm.locomotion.velocity.tail.speed_Mean' | -0,3580 | -0,4058 |
| 'worm.posture.kinks_Mean' | 0,0770 | 0,0729 | 'forward worm.locomotion.velocity.head.speed_Mean' | -0,3590 | -0,3832 |
| 'worm.posture.eigenProjection5_Q05' | 0,0552 | -0,0081 | 'absolute worm.locomotion.velocity.head.speed_Mean' | -0,3641 | -0,3926 |
| 'absolute worm.locomotion.bends.tail.amplitude_Q05' | 0,0541 | 0,0514 | 'worm.posture.amplitude.ratio_Q05' | -0,3656 | -0,3515 |
| 'worm.morphology.widthPerLength_Q95' | 0,0471 | 0,0709 | 'backward worm.locomotion.velocity.head.speed_Mean' | -0,3685 | -0,3993 |
| 'worm.posture.eccentricity_Q95' | 0,0432 | 0,0012 | 'absolute worm.locomotion.velocity.tailTip.speed_Mean' | -0,3686 | -0,4031 |
| 'absolute worm.posture.bends.neck.stdDev_Q05' | 0,0404 | 0,0468 | 'backward worm.locomotion.velocity.tailTip.speed_Q95' | -0,3694 | -0,4068 |
| 'absolute worm.posture.directions.head_Mean' | 0,0400 | 0,0464 | 'forward worm.locomotion.velocity.tailTip.speed_Mean' | -0,3847 | -0,4110 |
| 'worm.posture.eccentricity_Mean' | 0,0351 | 0,0209 | 'absolute worm.locomotion.bends.midbody.amplitude_Q05' | -0,3848 | -0,3723 |
| 'worm.posture.eigenProjection4_Mean' | 0,0273 | 0,0596 | 'worm.path.range_Q05' | -0,3898 | -0,4140 |
| 'absolute worm.posture.bends.head.mean_Q05' | 0,0187 | 0,0314 | 'absolute worm.posture.bends.hips.mean_Q95' | -0,3900 | -0,3267 |
| 'worm.posture.eigenProjection3_Mean' | 0,0105 | -0,0024 | 'absolute worm.locomotion.velocity.tailTip.speed_Q95' | -0,3941 | -0,4188 |

| | | | | | |
|---|---|---|---|---|---|
| 'worm.posture.eccentricity_Q05' | 0,0059 | 0,0018 | 'backward worm.locomotion.velocity.headTip.speed_Q95' | -0,3948 | -0,4030 |
| 'absolute worm.locomotion.bends.tail.frequency_Q05' | 0,0053 | -0,0040 | 'forward worm.locomotion.velocity.head.speed_Q95' | -0,3955 | -0,4007 |
| 'worm.posture.eigenProjection5_Mean' | -0,0059 | -0,0629 | 'absolute worm.locomotion.velocity.tail.speed_Mean' | -0,3965 | -0,4337 |
| 'absolute worm.locomotion.bends.tail.amplitude_Mean' | -0,0125 | -0,0252 | 'forward worm.locomotion.velocity.tailTip.speed_Q95' | -0,4001 | -0,4141 |
| 'absolute worm.locomotion.bends.head.frequency_Q95' | -0,0148 | -0,0207 | 'worm.posture.amplitude.ratio_Mean' | -0,4027 | -0,3980 |
| 'worm.posture.eigenProjection1_Mean' | -0,0161 | -0,0104 | 'absolute worm.locomotion.velocity.midbody.speed_Mean' | -0,4032 | -0,4350 |
| 'absolute worm.posture.bends.tail.mean_Q95' | -0,0196 | 0,0664 | 'backward worm.locomotion.velocity.tail.speed_Q95' | -0,4071 | -0,4455 |
| 'worm.posture.eigenProjection1_Q95' | -0,0272 | -0,0318 | 'forward worm.locomotion.velocity.tail.speed_Mean' | -0,4077 | -0,4379 |
| 'absolute worm.posture.directions.tail_Mean' | -0,0281 | -0,0458 | 'forward worm.locomotion.velocity.midbody.speed_Mean' | -0,4110 | -0,4360 |
| 'absolute worm.posture.bends.tail.stdDev_Q05' | -0,0353 | 0,0132 | 'absolute worm.locomotion.velocity.head.speed_Q95' | -0,4128 | -0,4229 |
| 'absolute worm.posture.bends.head.mean_Q95' | -0,0363 | 0,0016 | 'backward worm.locomotion.velocity.midbody.speed_Q95' | -0,4197 | -0,4509 |
| 'worm.posture.eigenProjection1_Q05' | -0,0450 | -0,0686 | 'absolute worm.locomotion.bends.head.amplitude_Mean' | -0,4276 | -0,3722 |
| 'worm.posture.eigenProjection5_Q95' | -0,0610 | -0,0853 | 'absolute worm.posture.bends.neck.mean_Mean' | -0,4315 | -0,3763 |
| 'absolute worm.locomotion.bends.tail.frequency_Q95' | -0,0636 | -0,0752 | 'backward worm.locomotion.velocity.head.speed_Q95' | -0,4335 | -0,4471 |
| 'absolute worm.locomotion.bends.tail.amplitude_Q95' | -0,0678 | -0,0562 | 'absolute worm.locomotion.velocity.tail.speed_Q95' | -0,4343 | -0,4585 |
| 'absolute worm.locomotion.bends.head.frequency_Q05' | -0,0869 | -0,0876 | 'forward worm.locomotion.velocity.tail.speed_Q95' | -0,4395 | -0,4529 |
| 'worm.posture.eigenProjection2_Mean' | -0,0886 | -0,1242 | 'worm.path.range_Mean' | -0,4426 | -0,4484 |
| 'backward worm.locomotion.velocity.headTip.speed_Q05' | -0,0923 | -0,1619 | 'worm.path.range_Q95' | -0,4452 | -0,4493 |
| 'absolute worm.posture.bends.head.stdDev_Q95' | -0,0994 | -0,0941 | 'absolute worm.locomotion.velocity.midbody.speed_Q95' | -0,4494 | -0,4695 |
| 'absolute worm.posture.bends.head.stdDev_Q05' | -0,1100 | -0,1406 | 'forward worm.locomotion.velocity.midbody.speed_Q95' | -0,4504 | -0,4610 |
| 'absolute worm.locomotion.bends.foraging.amplitude_Q05' | -0,1130 | -0,1533 | 'absolute worm.posture.bends.midbody.stdDev_Q05' | -0,5380 | -0,5202 |
| 'absolute worm.posture.bends.tail.stdDev_Q95' | -0,1172 | -0,0904 | 'absolute worm.locomotion.bends.midbody.amplitude_Q95' | -0,5657 | -0,5526 |
| 'backward worm.locomotion.velocity.tailTip.speed_Q05' | -0,1226 | -0,1804 | 'absolute worm.posture.bends.midbody.stdDev_Q95' | -0,5758 | -0,5289 |
| 'backward worm.locomotion.velocity.midbody.speed_Q05' | -0,1237 | -0,1692 | 'absolute worm.locomotion.bends.midbody.amplitude_Mean' | -0,6071 | -0,5989 |
| 'worm.posture.eigenProjection4_Q95' | -0,1249 | -0,0732 | 'absolute worm.posture.bends.midbody.stdDev_Mean' | -0,6314 | -0,5883 |
| 'backward worm.locomotion.velocity.tail.speed_Q05' | -0,1259 | -0,1786 | | | |
